# Extracting Knowledge from Scientific Texts on Patient-Derived Cancer Models Using Large Language Models: Algorithm Development and Validation

**DOI:** 10.1101/2025.01.28.634527

**Authors:** Jiarui Yao, Zinaida Perova, Tushar Mandloi, Elizabeth Lewis, Helen Parkinson, Guergana Savova

**Affiliations:** Computational Health Informatics Program, Boston Children’s Hospital, Harvard Medical School, 401 Park Drive, Boston, MA 02115, USA; European Molecular Biology Laboratory - European Bioinformatics Institute, Wellcome Genome Campus, Hinxton, Cambridge CB10 1SD, UK

**Author notes:** Corresponding author: Jiarui Yao. co-first authors (Jiarui Yao, Zinaida Perova).

## Abstract

Patient-derived cancer models (PDCMs) have emerged as indispensable tools in both cancer research and preclinical studies. The number of publications on PDCMs increased significantly in the last decade. Developments in Artificial Intelligence (AI), particularly Large Language Models (LLMs), hold promise for extracting knowledge from scientific texts at scale. This study investigates the use of LLM-based systems for automatically extracting PDCM-related entities from scientific texts. We evaluated two approaches: direct prompting and soft prompting using LLMs. For direct prompting, we manually create prompts to guide the LLMs to output PDCM-related entities from texts. The prompt consists of an instruction, definitions of entity types, gold examples and a query. We automatically train soft prompts – a novel line of research in this domain - as continuous vectors using machine learning approaches. Our experiments utilized state-of-the-art LLMs – proprietary GPT4-o and a series of open LLaMA3 family models. In our experiments, GPT4-o with direct prompts maintained competitive results. Our results demonstrate that soft prompting can effectively enhance the capabilities of smaller open LLMs, achieving results comparable to proprietary models. These findings highlight the potential of LLMs in domain-specific text extraction tasks and emphasize the importance of tailoring approaches to the task and model characteristics.

## Introduction

Patient-derived cancer models (PDCMs) are created from a patient’s own tumour sample and capture the complexity of human tumors to enable more accurate, personalized drug development and treatment selection. These models, including patient-derived xenografts (PDXs), organoids and cell lines, allow researchers to test treatments and identify most effective therapies, and have emerged as indispensable tools in both cancer research and precision medicine. The United States National Institutes of Health (NIH) has made significant investments in the generation and characterization of these models, with over 3 billion USD dedicated to active grants referencing PDCMs with a component of their research based on data extracted from the NIH Reporter [1] for FY 2024 alone. The number of publications using PDCMs continues to increase generating a substantial and rich metadata and data that needs standardization, harmonization and integration to maximize the impact of these models and their associated data within the research and clinical community. CancerModels.Org platform [2] serves as a unified gateway to the largest collection of PDCMs and related data. It empowers researchers and clinicians to discover suitable models for testing research hypotheses, conducting large-scale drug screenings, and advancing precision medicine initiatives. Extraction of PDCM-relevant knowledge and its harmonization within CancerModels.Org is essential to ensure that basic and translational researchers, bioinformaticians and tool developers have access to PDCM knowledge. While manual curation of publications ensures high accuracy when performed by domain experts, it is time-consuming and labor-intensive. Thus, a more streamlined and efficient knowledge acquisition method is needed to cater to the growing need of the scientific community for the timely availability of the PDCM metadata and associated data.

Large Language Models (LLMs) [3-5] often referred to as generative Artificial Intelligence (AI) systems are trained on vast amounts of data and have demonstrated impressive capabilities in the healthcare domain [6-8]. Researchers have studied LLMs in addressing various task related to healthcare such as diagnosing conditions [9-10], clinical decision support [11], answering patient questions [12], and medical education [13-14]. It has been shown that LLMs can extract meaningful information from texts [15-17].

In this work, we explore prompting techniques for LLMs with the goal to extract knowledge from PDCM-relevant scholarly publications. We applied two prompting methods – direct prompting [4] and soft prompting [18] with state-of-the-art (SOTA) proprietary and open LLMs. Our experimental results provide insights into choosing the optimal prompting methods for a specific task. The contributions of this paper are:

1. Studying the feasibility of SOTA LLMs as oncology knowledge extractors of PDCM-relevant information from scholarly scientific literature.
2. Creating a dataset of 100 abstracts of PDCM-relevant papers with entities annotated by experts.
3. Researching and comparing to our knowledge for the first time, direct vs. soft prompting techniques for oncology knowledge extraction, specifically PDCM-relevant information from scholarly scientific literature.

## Methods

### Concepts

We defined “knowledge” as entities of interest to researchers working with PDCMs within the oncology and cancer research fields. For example, the patient’s diagnosis provides a reference point to confirm that PDCM faithfully recapitulates the biology of the original tumour and is essential for ensuring the model’s relevance and reliability in studies of cancer progression or treatment response. Thus, “diagnosis” is important to understand the model’s characteristics in the context of disease in the patient. Patient’s age can significantly affect the molecular and genetic characteristics of the tumour. For example, pediatric cancers often have distinct genetic drivers and tumour microenvironments compared to cancers in older adults. In addition, age-related biological factors, such as immune system, metabolism and hormonal levels, influence how a tumour responds to treatments. Thus, knowing patient’s age is imperative for predictive accuracy of the model in preclinical testing and relevance of research findings.

We selected 15 most commonly used CancerModels.Org data model attributes (see Table 1), these include the attributes defined in the minimal information (MI) standard for patient-derived xenograft models (PDX-MI) [19] and draft MI standard for in vitro models [20].

**Table 1.** Entity definitions according to CancerModels.Org data model with examples and Inter-Annotator Agreement (IAA) F1 scores in the exact match setting which requires the spans of the annotators to match exactly.

| Entity type | Definition | Example | IAA |
| --- | --- | --- | --- |
| diagnosis | Diagnosis at the time of collection of the patient tumour used in the cancer model | triple-negative breast cancer, TNBC | 61.67 |
| age_category | Age category of the patient at the time of tissue sampling | adult, pediatric | 60.0 |
| genetic_effect | Any form of chromosomal rearrangement or gene-level changes | missense, amplification | 57.67 |
| model_type | Type of patient-derived model | PDX, organoid | 53.33 |
| molecular_char | Data or assay generated from or performed on the model in this study | RNA sequencing, whole-exome sequencing | 54.33 |
| biomarker | Gene, protein or biological molecule identified in or associated with patient's/model's tumour | BRCA1, IDH, epidermal growth factor receptor 2 | 61.33 |
| treatment | Treatment received by the patient or tested on the model | surgery, chemotherapy, PARP-inhibitor | 55.67 |
| response_to_treatment | Effect of the treatment on the patient's tumour or model | progression-free survival, reduced tumour growth | 55.0 |
| sample_type | The type of material used to generate the model or how this material was obtained | tissue fragment, autopsy | 49.0 |
| tumour_type | Collected tumour type used for generating the model | primary, recurrent | 49.67 |
| cancer_grade | Quantitative or qualitative grade reflecting how quickly the cancer is likely to grow | grade 1, low-grade | 42.0 |
| cancer_stage | Information about the cancer's extent in the body according to specific type of cancer staging system | TNM system, T0, stage I | 59.33 |
| clinical_trial | The type of clinical trial or Clinicaltrials.org identifier | phase II, prospective randomized clinical trials | 60.67 |
| host_strain | The name of the mouse host strain where the tissue sample was engrafted for generating the PDX model | NOD-SCID | 61.67 |
| model_id | ID of the patient-derived cancer model generated in this study | PHLC402 | 100.0 |

### Corpus

We used 100 abstracts as the gold standard corpus. The abstracts were chosen from the publications linked to the PDCMs submitted to CancerModels.Org platform and were selected to cover all three types of models in the resource - PDXs, organoids and cell lines. The final corpus is available on Github (see Data and code availability).

Three annotators (ZP, TM, EL) independently labelled entities in all 100 abstracts. It took 3 annotators about 40 hours in total to annotate 100 abstracts. The annotation quality was tracked through Inter-Annotator Agreement (IAA) - a measure of agreement between each annotation produced by different annotators working on the same dataset. The IAA is an indication of how difficult the task is for humans and it becomes the target for system development. We used pairwise F1 as a metric of the IAA [21] in the exact match setting which requires the spans of the annotators to match exactly. We computed the agreement between each pair of annotators and averaged across the three sets of scores. The final IAA F1 for each entity type is reported in Table 1. The final gold standard data were obtained through an adjudication step, where the annotators discussed the disagreements in their annotations and made the final decisions jointly. This dataset was then split into training, development (dev) and test sets in the standard 60:20:20 ratio. The train set was used for creating entity extraction algorithms, the development set – for refining the algorithms, and the test set – for the final evaluation. The number of instances of each entity type in each data split can be found in Table 2.

**Table 2.** Instances of each entity type in train, dev (development) and test sets respectively.

| Entity Type | Train | Dev | Test | TOTAL |
| --- | --- | --- | --- | --- |
| diagnosis | 362 | 122 | 114 | 598 |
| age_category | 19 | 0 | 0 | 19 |
| genetic_effect | 69 | 20 | 33 | 122 |
| model_type | 326 | 114 | 110 | 550 |
| molecular_char | 128 | 37 | 46 | 211 |
| biomarker | 503 | 118 | 163 | 784 |
| treatment | 426 | 77 | 130 | 633 |
| response_to_treatment | 99 | 21 | 28 | 148 |
| sample_type | 22 | 8 | 7 | 37 |
| tumour_type | 61 | 19 | 28 | 108 |
| cancer_grade | 6 | 1 | 1 | 8 |
| cancer_stage | 7 | 1 | 4 | 12 |
| clinical_trial | 35 | 2 | 4 | 41 |
| host_strain | 9 | 0 | 7 | 16 |
| model_id | 17 | 2 | 7 | 26 |
| TOTAL | 2,089 | 542 | 682 | 3,313 |

### Prompting Methods

Various prompting techniques have been proposed since the emergence of LLMs [22-25]. At a high level, these prompting techniques can be divided into two categories, direct prompting [4] and soft prompting [18,24,26]. The main difference between the two methods is the prompt representation, that is whether the prompt consists of human language words or vectors. Figure 1 shows the comparison of the two prompting methods. *Direct prompting* (or discrete prompting) is the most intuitive prompting method where users directly interact with LLMs using natural language. For example, a user may ask ChatGPT to “Write a thank you note to an old friend of my parents”, in this case the text within the quotation marks is a discrete prompt. *Soft prompting* (or continuous prompting) uses a machine learning approach to train a sequence of continuous vectors, which are the “virtual tokens” of the prompt. It is worth noting that soft prompting differs from finetuning. With soft prompting, the parameters in the LLM are not updated, only the parameters in the soft prompt are updated. In contrast, finetuning requires to update the parameters in the entire LLM, therefore needs more computation resource. Both prompting techniques have their advantages and disadvantages. Compared to direct prompting, soft prompting does not require the tedious process of manually creating prompts, however, it requires some labeled data to train the prompt. In this work, we explore both direct and soft prompting as we aim to explore the latest developments in LLMs and prompting techniques for the task of extracting PDCM entities from abstracts of academic papers.

**Figure 1.**
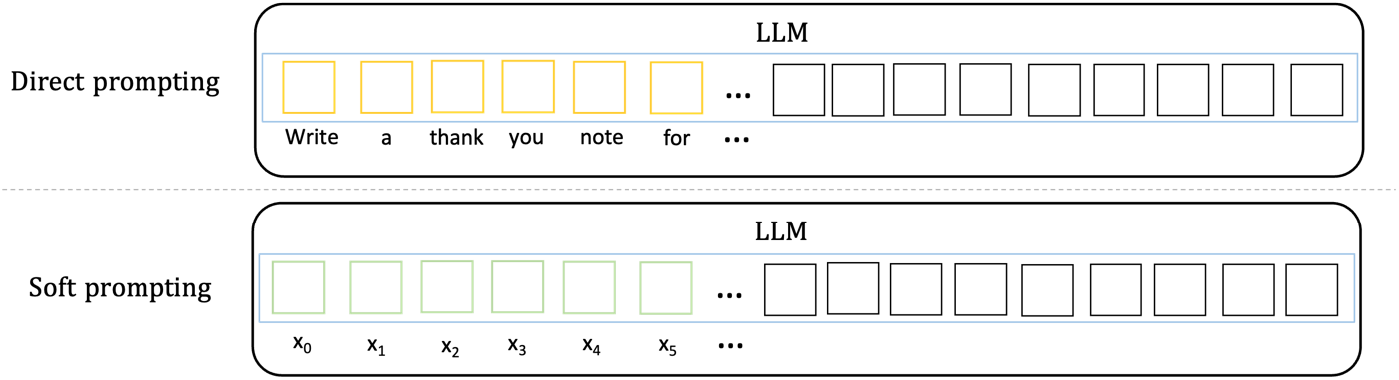
Illustration of the two prompting methods. In direct prompting, a prompt contains a sequence of words. In soft prompting, a prompt consists of a list of vectors.

### Direct Prompting

When asking LLMs to extract entities such as diagnoses or biomarkers, the most intuitive way is to ask LLMs to output the entities directly. In Example 1 below, “ALK” is a biomarker entity. One may expect the model to output *{“biomarker”: [ALK]}*. However, we note that the string “ALK” is mentioned multiple times in this example text, therefore it is not clear which “ALK” the model refers to. To get the most precise extraction to facilitate a more fine-grained analysis, we instruct the model to output the offsets of the specific mentions in the text (i.e. the spans). For instance, if the model gives us *[(48, 51, “ALK”, biomarker), (323, 326, “ALK”, biomarker), …]*, we know that from character 48 to character 51, there is a biomarker entity, “ALK”. Similarly, we can find another biomarker entity “ALK” at position 323 – 326.

#### Example 1

*“Oncogenic fusion of anaplastic lymphoma kinase (ALK) with echinoderm microtubule associated protein like 4 protein or other partner genes occurs in 3% to 6% of lung adenocarcinomas. Although fluorescence in situ hybridization (FISH) is the accepted standard for detecting anaplastic lymphoma receptor tyrosine kinase gene (ALK) gene rearrangement that gives rise to new fusion genes, not all ALK FISH-positive patients respond to ALK inhibitor therapies*.*”*

We started our exploration by designing prompts that included an explicit instruction to specify the character offsets of each entity along with the entity text and type (e.g., *48, 51, “ALK”, biomarker*). However, our experiments show that it was challenging for the LLM to output the correct character offsets, a seemingly straightforward task (all the model needs to do is to count the number of characters); however the complexity of this seemingly straightforward task is likely due to the LLM’s way of breaking words outside its vocabulary into so-called word pieces, e.g. “organoid” is broken down into two word pieces “organ” and “-oid”. Considering that LLMs are trained as generative models [3-4], we subsequently cast the entity extraction task as a generation task, where we instruct the model to mark the entities with XML tags. For instance, if the model outputs “Oncogenic fusion of anaplastic lymphoma kinase (<biomarker>ALK</biomarker>) with echinoderm microtubule …”, then postprocessing the output with regular expressions would find the exact position of “ALK” in the text. Specifically, we ask the LLMs to mark the start and end of an entity with <entity_type> and </entity_type> tags, where entity_type is a placeholder for the specific entity type, such as biomarker or treatment (see Table 1 for the full list).

### Soft Prompting

Designing the direct prompts manually could be time-consuming, especially when using the technique of chain-of-thought (CoT) prompting [22] as it requires human input for the step-by-step reasoning examples for the LLMs [27]. In addition, minor changes in the prompt language could lead to drastic changes in the model performance [24]. On the other hand, soft prompting requires some amount of gold data to be used for its training and annotating gold data by domain experts could also be time-consuming. Fortunately, only a small set of labeled data is needed to train soft prompts. As pointed out above, we labeled 100 abstracts, which we utilize for training and evaluating the soft prompts.

There are a few soft prompting methods, the difference usually lies in how the prompt vectors are initialized and learned. Prompt-tuning [18] is a technique that learns the prompt by adding a list of virtual tokens (i.e., vectors) in front of the input, where the virtual tokens can be randomly initialized, or drawn from a pre-trained word embedding [28] set. Another method is P-tuning [24], which employs small neural networks such as feedforward neural networks [29] (multilayer perceptron or MLP) or recurrent neural networks [30] (e.g., long-short term memory or LSTM) as prompt encoder to learn the prompt. Only the parameters in the prompt encoder are updated during training, while the weights in the LLMs are frozen. In our experiments, we found P-tuning did not always converge to the optimal solution for our task perhaps due to the random initialization of the vectors rather than using carefully pre-trained word embeddings. Therefore, we focus on prompt-tuning in this work. Following Lester et al. [18], we initialize the vectors in the prompt with the embeddings of the labels in the entity type set (see Table 1).

The standard approach for entity extraction in natural language processing (NLP) is via token classification [31]. Concretely, a classifier is trained to predict the label for each token in a sentence according to a pre-defined label set. Additionally, each label is prepended with a B or I prefix to indicate the entity’s **b**eginning or **i**nside mention, respectively. An example is provided in Table 3. “Ewing sarcoma” is an entity mention of diagnosis type. Thus “Ewing” and “sarcoma” are labeled as “Diagnosis”, while all other tokens are labeled as “O”, meaning they are **O**utside of an entity. To be more precise, “Ewing” is at the beginning of the diagnosis entity, and “sarcoma” is inside of the entity, so they are labeled as “**B**-Diagnosis” and “**I**-Diagnosis” respectively.

**Table 3.** An example of entity extraction as token classification.

|  |  |  |  |  |  |  |  |  |  |  |
| --- | --- | --- | --- | --- | --- | --- | --- | --- | --- | --- |
| Ewing | sarcoma | is | a | tumour | of | the | bone | and | soft | tissue |
| B-Diagnosis | I-Diagnosis | O | O | O | O | O | O | O | O | O |

To summarize, we train a multi-class classifier for the soft-prompting training step. There are 15 entity types in our dataset, therefore there are 15 * 2 + 1 = 31 labels for token classification, with one extra label for “O”.

### Experimental Set-up

For efficiency purposes, we use Apache cTAKES [32] to split an abstract into sentences which are then passed to the LLMs to extract entities one sentence at a time. Our direct prompt includes the instruction, definition of each entity, five examples (few-shot In-Context Learning (ICL)) and the query (the sentence). The ICL [4] is a common practice in LLM prompting and has consistently shown improved results as the examples guide the LLM onto an optimal path [33-34]. Figure 2 presents the prompt template, examples are in Supplementary files Table 1.

**Figure 2.**
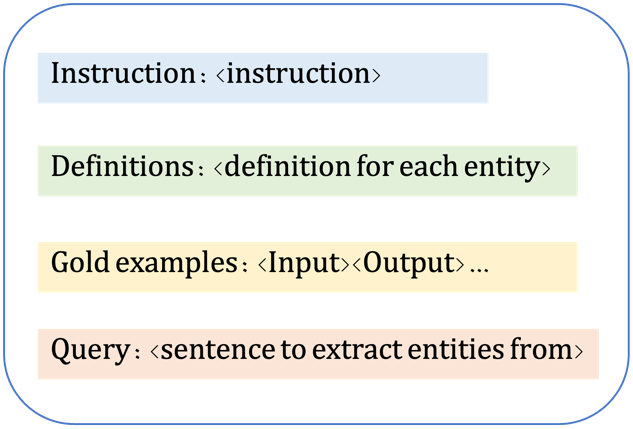
: Prompt template used in direct prompting.

When choosing the LLMs, we used GPT-4o [35], one of the most powerful proprietary LLMs at the time of this study, and SOTA open LLMs from the LLaMA3 family [36], including LLaMA3.1 70B, LLaMA3.1 8B, LLaMA3.2 1B and LLaMA3.2 3B. We did not use GPT-4o or LLaMA3.1 70B to train the soft prompts due to computational resource limitations; thus our work here is representative of the computational environment in the vast majority of academic medical centers and research labs. We set the soft prompt length to 30. We trained the soft prompt on the training set for 50 epochs with a learning rate of 0.001. Hyper-parameters were tuned on the development set using the LLaMA3.1 8B model. We report the evaluation results on the test set in the next section.

## Results

We use the standard evaluation metrics of precision/positive predictive value, recall/sensitivity and F1 (the harmonic mean of precision and recall). We consider two evaluation settings: “exact match” setting requires the span output from the model to exactly match the span of the gold annotation, and “overlapping match” setting allows the model to get partial credit if its extraction overlaps the spans in the gold annotation. For example, the model may extract “patient-derived tumor xenograft (PDX)” as a model_type entity, while the gold annotation is “patient-derived tumor xenograft (PDX) **models**”. Under the “exact match” setting, “patient-derived tumor xenograft (PDX)” is NOT a match to “patient-derived tumor xenograft (PDX) **models**”; while under the “overlapping match” setting, they are a match since the spans overlap.

Tables 4 and 5 show the evaluation results. In Table 4, we can see that under the “exact match” setting, direct prompting GPT-4o achieves the highest F1, 50.48. The performances of LLaMA3 family models drop as the model size decreases, from 38.40 F1 of the 70B model to 6.78 F1 of the 1B model. Despite the suboptimal performance of direct prompting of the smaller LLaMA3 models, we see a consistent improvement in F1 scores when using soft prompting. Especially for the LLaMA3.2 models, the performance of the 3B model increases from 7.06 F1 to 46.68 F1, over 8 points higher than the LLaMA3.1-70B model with direct prompting (38.40 F1), despite the huge disparities in their model size.

**Table 4:** Evaluation results (exact match) as precision/positive predictive value, recall/sensitivity and F1 (harmonic mean of precision and recall). Best results are in bold.

| Exact match | Precision | Recall | F1 |
| --- | --- | --- | --- |
| Direct prompting |  |  |  |
| GPT-4o | 56.09 | **45.89** | **50.48** |
| LLaMA3.1-70B | **57.27** | 28.89 | 38.40 |
| LLaMA3.1-8B | 35.80 | 18.48 | 24.37 |
| LLaMA3.2-3B | 25.23 | 4.10 | 7.06 |
| LLaMA3.2-1B | 23.48 | 3.96 | 6.78 |
| Soft prompting |  |  |  |
| LLaMA3.1-8B | 47.17 | 45.75 | 46.44 |
| LLaMA3.2-3B | **47.30** | **46.09** | **46.68** |
| LLaMA3.2-1B | 46.19 | 45.01 | 45.59 |

**Table 5:** Evaluation results (overlapping match) as precision/positive predictive value, recall/sensitivity and F1 (harmonic mean of precision and recall). Best results are in bold.

| Overlapping match | Precision | Recall | F1 |
| --- | --- | --- | --- |
| Direct prompting |  |  |  |
| GPT-4o | 76.96 | **66.52** | **71.36** |
| LLaMA3.1-70B | **77.95** | 43.99 | 56.24 |
| LLaMA3.1-8B | 50.54 | 27.49 | 35.61 |
| LLaMA3.2-3B | 41.03 | 7.03 | 12.0 |
| LLaMA3.2-1B | 35.34 | 6.01 | 10.28 |
| Soft prompting |  |  |  |
| LLaMA3.1-8B | 71.19 | 70.53 | 70.86 |
| LLaMA3.2-3B | **72.05** | **71.55** | **71.80** |
| LLaMA3.2-1B | 70.38 | 70.48 | 70.42 |

Similar trends are observed in Table 5 under the “overlapping match” evaluation. GPT4-o shows a performance of 71.36 F1, remaining firmly at the top position using direct prompting. The three smaller LLaMA3 models continue to benefit from soft prompting, with the LLaMA3.2 3B model achieving slightly higher score than GPT4-o with direct prompting (71.80 F1 vs. 71.36 F1).

## Discussion

Our experiments show that soft prompting can dramatically boost the performance of smaller LLMs. The performance of the three LLaMA models with soft prompting are similar, about 46 F1 in the exact match setting, and 70’s F1 in the overlapping match setting. These are promising results given the limited number of abstracts used for training - 60 abstracts with 2,089 entity mentions.

How much data is needed to train the soft prompt? To answer this question, we trained LLaMA3.2 1B model, the smallest model used in this work, with different amounts of training data. Figure 3 shows the relation between the proportion of training data and the F1 scores on the test set (overlapping match). We can see that solid performance was achieved with only 5% of the training data, which is 26 sentences (about 3 abstracts). With 25% of the training data, i.e. 129 sentences (about 15 abstracts), the model obtained 68.21 F1 score, only 2 points lower than using the whole training set, and only 3 points lower than GPT4-o with direct prompting. Despite the impressive performance of direct prompting GPT4-o, one potential issue is that not all data used in biomedical research can be sent to proprietary models such as GPT or Gemini family models [8] via public APIs. That is, for applications that use real patient data which require HIPPA-compliant platforms, our work shows that it is still possible to reach performance close to proprietary LLMs such as GPT4-o by training soft prompts, with a tradeoff for the need of a small set of labeled data.

**Figure 3.**
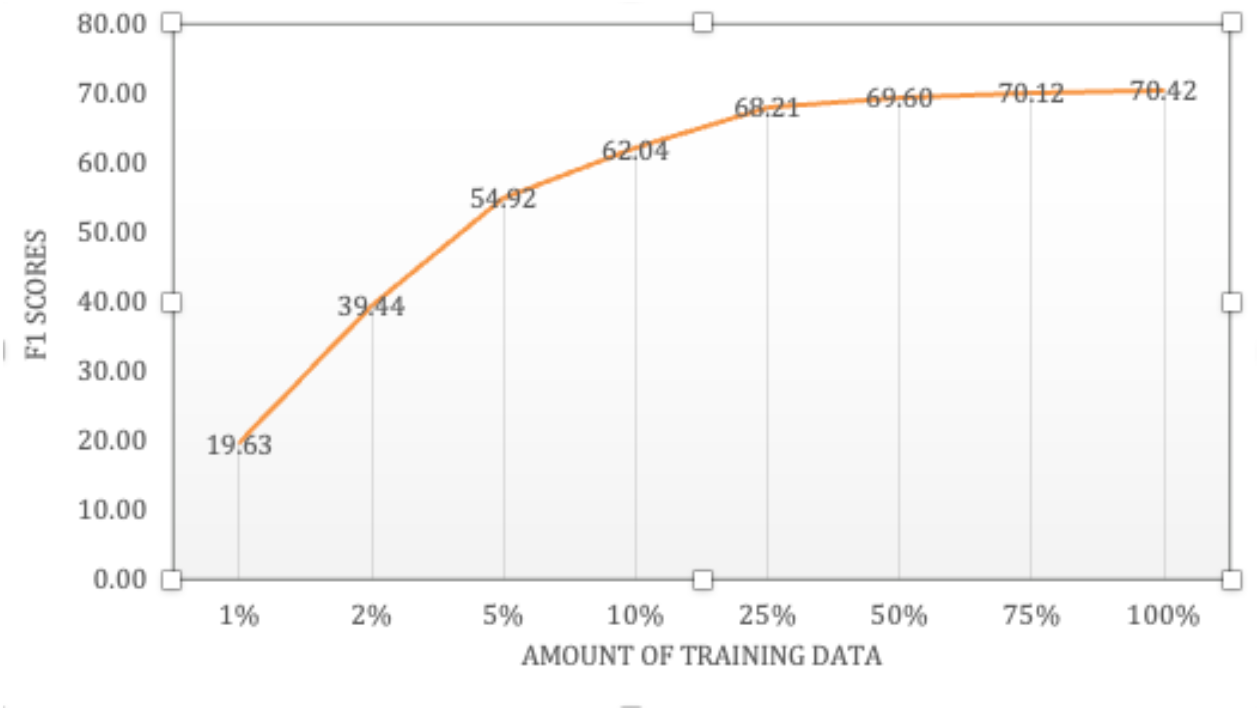
Model performances of the LLaMA3.2 1B model as the size of training data increases.

Some entities appear more frequently than other entities in our data. For example, diagnosis and treatment are mentioned more often than entities such as cancer grade. In Table 6, we present the number of instances of each entity type in our dataset and the corresponding GPT4-o with direct prompting performance in overlapping setting for that entity. We can see that GPT4 performs the best for the entity types that have the most instances -- diagnosis, model type, treatment entities. Of these frequent entity types, biomarker is the one with the lowest performance. Several factors could have contributed to these results, including ambiguous and inconsistent mentions and contextual dependencies. In this task, we defined biomarker as “gene, protein or biological molecule identified in or associated with patient’s/model’s tumour.” Thus, biomarker entities can be mentioned using their full names (e.g., epidermal growth factor receptor, lnc-RP11-536 K7.3, echinoderm microtubule-associated protein-like 4), standardized gene or protein symbols (*NPM1*, KRAS, PTEN) or abbreviation of metabolites (NADPH, D2HG). Moreover, biomarker entity (e.g., “MEK”) often overlaps with treatment entity (e.g., “MEK inhibitor”). The ambiguity in biomarker entity mentions might interfere with the model’s ability to recognize them consistently. In addition, biomarker entities are often mentioned as lists (see Example 2) resulting in a different frequency within and across the abstracts and patterns of entity mentions, in comparison with other entities.

**Table 6:** Evaluation results of GPT4-o with direct prompts for each entity type as precision/positive predictive value, recall/sensitivity and F1 (harmonic mean of precision and recall). Results are overlapping match setting on the test set. Top F1 scores are in bold. F1 scores exceeding the IAA are underlined.

| Entity Type | # Instances Train | # Instances Dev | # Instances Test | Precision | Recall | F1 | IAA |
| --- | --- | --- | --- | --- | --- | --- | --- |
| diagnosis | 362 | 122 | 114 | 92.47 | 75.44 | **<u>83.09</u>** | 61.67 |
| age_category | 19 | 0 | 0 | 0.0 | 0.0 | 0.0 | 60.0 |
| genetic_effect | 69 | 20 | 33 | 45.71 | 47.06 | 46.38 | 57.67 |
| model_type | 326 | 114 | 110 | 88.07 | 84.21 | **<u>86.10</u>** | 53.33 |
| molecular_char | 128 | 37 | 46 | 65.22 | 63.83 | <u>64.52</u> | 54.33 |
| biomarker | 503 | 118 | 163 | 85.05 | 55.49 | <u>67.16</u> | 61.33 |
| treatment | 426 | 77 | 130 | 81.74 | 70.15 | **<u>75.50</u>** | 55.67 |
| response_to_treatment | 99 | 21 | 28 | 38.64 | 60.71 | 47.22 | 55.0 |
| sample_type | 22 | 8 | 7 | 45.45 | 71.43 | <u>55.56</u> | 49.0 |
| tumour_type | 61 | 19 | 28 | 66.67 | 57.14 | <u>61.54</u> | 49.67 |
| cancer_grade | 6 | 1 | 1 | 50.0 | 100 | <u>66.67</u> | 42.0 |
| cancer_stage | 7 | 1 | 4 | 33.33 | 25.0 | 28.57 | 59.33 |
| clinical_trial | 35 | 2 | 4 | 80.0 | 100 | **<u>88.89</u>** | 60.67 |
| host_strain | 9 | 0 | 7 | 100 | 28.57 | 44.44 | 61.67 |
| model_id | 17 | 2 | 7 | 66.67 | 28.57 | 40.0 | 100.0 |

### Example 2

*Genomic alterations involved RB1 (55%), TP53 (46%), PTEN (29%), BRCA2 (29%), and AR (27%), and there was a range of androgen receptor signaling and NEPC marker expression*.

The extraction of PDCM-relevant knowledge is not an easy task for the domain experts as indicated by the IAA which below 65 F1 for all entity types except for model_id. In 9 out of 15 entity types, the system performance in an overlapping match setting exceeds substantially the IAA (last two columns of Table 6). This is the case for categories with plentiful training instances (e.g. diagnosis, model_type) as well as for categories with fewer training instances (e.g. sample_type, tumour_type). For the exact match setting, in 6 out of 15 entity types, the system performance exceeds the IAA (last two columns in Table 7). Therefore, the LLM could be a feasible assistant whose output is reviewed by a domain expert to ascertain the final abstraction. We think such human-in-the-loop scenarios are a fruitful path forward.

**Table 7:** Evaluation results of GPT4-o with direct prompts for each entity type as precision/positive predictive value, recall/sensitivity and F1 (harmonic mean of precision and recall). Results are exact match setting on the test set. Top F1 scores are in bold. F1 scores exceeding the IAA are underlined.

| Entity Type | # Instances Train | # Instances Dev | # Instances Test | Precision | Recall | F1 | IAA |
| --- | --- | --- | --- | --- | --- | --- | --- |
| diagnosis | 362 | 122 | 114 | 77.17 | 62.28 | **<u>68.93</u>** | 61.67 |
| age_category | 19 | 0 | 0 | 0.0 | 0.0 | 0.0 | 60.0 |
| genetic_effect | 69 | 20 | 33 | 25.71 | 27.27 | 26.47 | 57.67 |
| model_type | 326 | 114 | 110 | 56.88 | 56.36 | **<u>56.62</u>** | 53.33 |
| molecular_char | 128 | 37 | 46 | 54.35 | 54.35 | **<u>54.35</u>** | 54.33 |
| biomarker | 503 | 118 | 163 | 46.74 | 26.38 | 33.73 | 61.33 |
| treatment | 426 | 77 | 130 | 72.34 | 52.31 | **<u>60.71</u>** | 55.67 |
| response_to_treatment | 99 | 21 | 28 | 27.91 | 42.86 | 33.80 | 55.0 |
| sample_type | 22 | 8 | 7 | 45.45 | 71.43 | **55.56** | 49.0 |
| tumour_type | 61 | 19 | 28 | 50.0 | 39.29 | 44.0 | 49.67 |
| cancer_grade | 6 | 1 | 1 | 50.0 | 100 | **66.67** | 42.0 |
| cancer_stage | 7 | 1 | 4 | 33.33 | 25.0 | 28.57 | 59.33 |
| clinical_trial | 35 | 2 | 4 | 40.0 | 50.0 | 44.44 | 60.67 |
| host_strain | 9 | 0 | 7 | 100 | 14.29 | 25.0 | 61.67 |
| model_id | 17 | 2 | 7 | 66.67 | 28.27 | 40.0 | 100.0 |

We would like to note that the work presented in the paper was done in a computational environment representative of the vast majority of academic medical centers and non-industry research labs. Although we have access to SOTA GPUs, we still find ourselves constrained as to the extent to which we could utilize very large language models and the research we could potentially execute. The larger community needs to address the growing gap of computational resources between big tech and the rest of the research community.

## Limitations

As this is a feasibility study, we limited ourselves to the extraction of entity mentions of 15 entity types and annotation of 100 abstracts. The entity types in this task were chosen from attributes in the descriptive standards for PDCMs. While these are recognised by the PDCM and oncology community, they do not cover all knowledge in the PDCM-relevant texts. Some refinement of the entity types will be beneficial to improve prompting results.

We limited our corpus to 100 abstracts from papers, associated with PDCMs deposited in CancerModels.Org. We did not assess the abstracts for the presence and/or equal distribution of all the entities. Thus, there were very few mentions of some of the entities in the corpus (e.g. cancer_stage), negatively affecting our overall F1. We decided not to exclude those entities, as these results can guide refinements of the future studies. The computational methods discussed here are applicable to other studies that require the extraction of textual information from scientific papers. Future work could involve extending this method to extract knowledge from the main body of the papers.

## Conclusions

This study highlights the promising potential of LLMs as powerful tools for extracting meaningful PDCM-relevant knowledge from scientific literature, a task critical to advancing cancer research and precision medicine. By comparing direct and soft prompting techniques with both proprietary and open LLMs, we provide insights into the most effective methods for knowledge extraction in PDCM-relevant oncology domain. We show that GPT4-o with direct prompts maintained competitive results and soft prompting helped improve the performance of smaller LLMs by a large margin. In conclusion, our work shows that it is possible to achieve the performance of proprietary LLMs by training soft prompts with smaller open models.

To our knowledge, this is the first study implementing SOTA LLMs in the PDCM domain for knowledge extraction and the first study to research the hot topic of soft prompting in this domain. Our results show that LLMs can streamline the acquisition of complex cancer model metadata. This can potentially reduce the burden of manual curation and speed up the integration of PDCMs into research and clinical workflows. This work also paves the way for future studies on optimizing LLMs for large-scale knowledge extraction tasks. Ultimately, the ability to efficiently extract and harmonise PDCM-relevant knowledge will help drive progress in cancer research and precision oncology, providing researchers and clinicians with better tools to improve patient’s outcomes.

At a higher level, our study contributes to the growing body of research into understanding what tasks benefit from using LLMs as LLMs are likely not the perfect technology that could solve every single task.

## Data and code availability

The data and code will be available upon publication in the CancerModels.Org Github repository [37].

## Acknowledgments

Funding was provided by US National Institutes of Health (U24CA248010, R01LM013486, U24CA253539) and European Bioinformatics Institute (EMBL-EBI) Core Funds.

## Supplementary files

Prompts used in direct prompting experiments are presented in the Supplementary files.

