## Supplemental Table 1 for "Extracting Knowledge from Scientific Texts on Patient-Derived Cancer Models Using Large Language Models: Algorithm Development and Validation"

Table 1. Instruction and examples used in direct prompting experiments.

| | |
|---|---|
| Instruction | You will be given a sentence from a paper on PDCM. Please extract entities as defined below and return as an XML. In the XML, please mark the start and end of the entity with <entity_type></entity_type>. Please return one entity at one time. That is, if there are n entities in the sentence, print the sentence with the marked entity for n times. Do not change anything else in the sentence. |
| Examples | Input:<br>There were 13 missense mutations identified in the xenograft that were not present in the patient's primary tumor and there were no new nonsense mutations.<br>Output:<br>The entities in this sentence are:<br>{"genetic_effect": [missense mutations], "model_type": [xenograft], "tumour_type": [primary]}<br>There are 3 entities in total. Now I will print out the sentence for 3 times, each time with only one entity marked, the rest of the sentence will be exactly the same as the original sentence.<br>(1) There were 13 <genetic_effect>missense mutations</genetic_effect> identified in the xenograft that were not present in the patient's primary tumor and there were no new nonsense mutations.<br>(2) There were 13 missense mutations identified in the <model_type>xenograft</model_type> that were not present in the patient's primary tumor and there were no new nonsense mutations.<br>(3) There were 13 missense mutations identified in the xenograft that were not present in the patient's <tumour_type>primary</tumour_type> tumor and there were no new nonsense mutations.<br><br>Input:<br>Here we present the detailed molecular, cellular, genetic and epigenetic characterization of an orthotopic xenograft derived from a high-risk stage 4 neuroblastoma patient.<br>Output:<br>The entities in this sentence are:<br>{"molecular_char": [molecular, cellular, genetic, epigenetic characterization], "model_type": [orthotopic xenograft], "cancer_stage": [stage 4], "diagnosis": [neuroblastoma]}<br>There are 7 entities in total. Now I will print out the sentence for 7 times, each time with only one entity marked, the rest of the sentence will be exactly the same as the original sentence.<br>(1) Here we present the detailed <molecular_char>molecular<molecular_char>, cellular, genetic and epigenetic characterization of an orthotopic xenograft derived from a high-risk stage 4 neuroblastoma patient.<br>(2) Here we present the detailed molecular, <molecular_char>cellular</molecular_char>, genetic and epigenetic characterization of an orthotopic xenograft derived from a high-risk stage 4 neuroblastoma patient.<br>(3) Here we present the detailed molecular, cellular, <molecular_char>genetic</molecular_char> and epigenetic characterization of an orthotopic xenograft derived from a high-risk stage 4 neuroblastoma patient. |

| | |
|---|---|
| | (4) Here we present the detailed molecular, cellular, genetic and <molecular_char>epigenetic characterization</molecular_char> of an orthotopic xenograft derived from a high-risk stage 4 neuroblastoma patient.<br>(5) Here we present the detailed molecular, cellular, genetic and epigenetic characterization of an <model_type>orthotopic xenograft</model_type> derived from a high-risk stage 4 neuroblastoma patient.<br>(6) Here we present the detailed molecular, cellular, genetic and epigenetic characterization of an orthotopic xenograft derived from a high-risk <cancer_stage>stage 4</cancer_stage> neuroblastoma patient.<br>(7) Here we present the detailed molecular, cellular, genetic and epigenetic characterization of an orthotopic xenograft derived from a high-risk stage 4 <diagnosis>neuroblastoma</diagnosis> patient.<br><br><br>Input:<br>Neuroblastoma is a pediatric cancer of the developing sympathoadrenal lineage.<br>Output:<br>The entities in this sentence are:<br>{"diagnosis": [Neuroblastoma], "age_category": [pediatric]}<br>There are 2 entities in total. Now I will print out the sentence for 2 times, each time with only one entity marked, the rest of the sentence will be exactly the same as the original sentence.<br>(1) <diagnosis>Neuroblastoma</diagnosis> is a pediatric cancer of the developing sympathoadrenal lineage.<br>(2) Neuroblastoma is a <age_category>pediatric</age_category> cancer of the developing sympathoadrenal lineage.<br><br><br>Input:<br>With these PDXs, we performed in vivo compound screens using a 1 × 1 × 1 experimental design (PDX clinical trial or PCT) to assess the population responses to 62 treatments across six indications.<br>Output:<br>The entities in this sentence are:<br>{"model_type": [PDXs], "clinical_trial": [PDX clinical trial, PCT]}<br>There are 3 entities in total. Now I will print out the sentence for 3 times, each time with only one entity marked, the rest of the sentence will be exactly the same as the original sentence.<br>(1) With these <model_type>PDXs<model_type>, we performed in vivo compound screens using a 1 × 1 × 1 experimental design (PDX clinical trial or PCT) to assess the population responses to 62 treatments across six indications.<br>(2) With these PDXs, we performed in vivo compound screens using a 1 × 1 × 1 experimental design (<clinical_trial>PDX clinical trial</clinical_trial> or PCT) to assess the population responses to 62 treatments across six indications.<br>(3) With these PDXs, we performed in vivo compound screens using a 1 × 1 × 1 experimental design (PDX clinical trial or <clinical_trial>PCT<clinical_trial>) to assess the population responses to 62 treatments across six indications.<br><br><br>Input: |

| | |
|---|---|
| | As autopsy specimens had an ALK R1181C mutation, PDX tumor bearing animals were treated with the pan-kinase inhibitor lestaurtinib but demonstrated no decrease in tumor growth.<br>Output:<br>The entities in this sentence are:<br>{"sample_type": [autopsy], "biomarker": [ALK], "model_type": [PDX], "treatment": [pan-kinase inhibitor, lestaurtinib], "response_to_treatment": [decrease in tumor growth]}<br>There are 6 entities in total. Now I will print out the sentence for 6 times, each time with only one entity marked, the rest of the sentence will be exactly the same as the original sentence.<br>(1) As <sample_type>autopsy</sample_type> specimens had an ALK R1181C mutation, PDX tumor bearing animals were treated with the pan-kinase inhibitor lestaurtinib but demonstrated no decrease in tumor growth.<br>(2) As autopsy specimens had an <biomarker>ALK<biomarker> R1181C mutation, PDX tumor bearing animals were treated with the pan-kinase inhibitor lestaurtinib but demonstrated no decrease in tumor growth.<br>(3) As autopsy specimens had an ALK R1181C mutation, <model_type>PDX</model_type> tumor bearing animals were treated with the pan-kinase inhibitor lestaurtinib but demonstrated no decrease in tumor growth.<br>(4) As autopsy specimens had an ALK R1181C mutation, PDX tumor bearing animals were treated with the <treatment>pan-kinase inhibitor</treatment> lestaurtinib but demonstrated no decrease in tumor growth.<br>(5) As autopsy specimens had an ALK R1181C mutation, PDX tumor bearing animals were treated with the pan-kinase inhibitor <treatment>lestaurtinib</treatment> but demonstrated no decrease in tumor growth.<br>(6) As autopsy specimens had an ALK R1181C mutation, PDX tumor bearing animals were treated with the pan-kinase inhibitor lestaurtinib but demonstrated no <response_to_treatment>decrease in tumor growth</response_to_treatment>. |
